# Induction and suppression of NF-_k_B signalling by a DNA virus of *Drosophila*

**DOI:** 10.1101/358176

**Authors:** William H. Palmer, Joep Joosten, Gijs J. Overheul, Pascal W. Jansen, Michiel Vermeulen, Darren J. Obbard, Ronald P. Van Rij

## Abstract

Interactions between the insect immune system and RNA viruses have been best studied in *Drosophila*, where RNA interference, NF-_k_B and JAK-STAT pathways underlie antiviral immunity. In response to these immune mechanisms, insect viruses have convergently evolved suppressors of RNA interference that act by diverse mechanisms to permit viral replication. However, interactions between the insect immune system and DNA viruses have received less attention, primarily because few *Drosophila-infecting* DNA virus isolates are available. Here, we use a recently-isolated DNA virus of *Drosophila melanogaster*, Kallithea virus, to probe known antiviral immune responses and virus evasion tactics in the context of DNA virus infection. We find that fly mutants for RNA interference and Immune deficiency (Imd), but not Toll, pathways are more susceptible to Kallithea virus infection. We identify the Kallithea virus-encoded protein gp83 as a potent inhibitor of Toll signalling, strongly suggesting that Toll mediates antiviral responses during Kallithea virus infection, but that it is suppressed by the virus. Further, we find that Kallithea gp83 inhibits Toll signalling either through NF-_k_B transcription factor regulation, or transcriptionally. Together, these results provide a broad description of known antiviral pathways in the context of DNA virus infection and identify the first Toll pathway inhibitor in a *Drosophila* virus, extending the known diversity of insect virus-encoded immune inhibitors.

## Introduction

Innate antiviral immunity in insects has been best studied in response to RNA virus infections of *Drosophila melanogaster.* Antiviral immune mechanisms that target RNA viruses include RNA-mediated defences such as RNA interference (RNAi) and RNA decay pathways, cellular defences such as apoptosis, phagocytosis, and autophagy, and transcriptional responses. The latter are primarily mediated by Janus kinase/signal transducers and activators of transcription (JAK-STAT) and Nuclear factor κB (NF-_k_B) pathways (reviewed in Merkling and van Rij, 2013; Bronkhorst and van Rij, 2014; Lamiable and Imler, 2014; Xu and Cherry, 2014; Palmer, Varghese and van Rij, 2018).

The insect response to DNA viruses is less well studied, but RNAi and apoptosis have demonstrated antiviral activity (Clem, 2001; Bronkhorst *et al.*, 2012; Kemp *et al.*, 2013) and the JAK-STAT pathway is active during infection, possibly mediating a tolerance response (West and Silverman, 2018). Baculovirus, nudivirus, and iridovirus infections of *Drosophila* all give rise to virus-derived small interfering RNA (vsiRNAs), which regulate DNA virus gene expression (Bronkhorst *et al.*, 2012; Jayachandran, Hussain and Asgari, 2012; Kemp *et al.*, 2013; Webster *et al.*, 2015) and mutants for RNAi effectors *Dicer-2 (Dcr-2)* and *Argonaute-2 (AGO2)* are hypersensitive to Invertebrate iridescent virus 6 (IIV6; an iridovirus) infection. This suggests that RNAi is also an important defence against DNA viruses, and IIV6 correspondingly encodes a suppressor of RNAi (Bronkhorst *et al.*, 2012, 2014). Virus-encoded suppressors of apoptosis are also widespread in DNA viruses, acting through binding and inhibition of cellular caspases (e.g. p35), or stabilization of cellular inhibitors of apoptosis (e.g. IAP gene family; Bump *et al.*, 1995; Xue and Robert Horvitz, 1995; Byers, Vandergaast and Friesen, 2016). In contrast, the contribution of transcriptional responses, such as the NF-_k_B pathways, to DNA viruses has not yet been elucidated.

There are two NF-_k_B pathways in *Drosophila:* Toll and Imd, which primarily function in antibacterial (Toll: gram-positive, Imd: gram-negative) and antifungal (Toll) defense, although both provide protection against some RNA viruses (reviewed in Valanne, Wang and Ramet, 2011; Merkling and van Rij, 2013; Lamiable and Imler, 2014; Myllymaki, Valanne and Ramet, 2014; Palmer, Varghese and van Rij, 2018). Toll and Imd pathways are activated following recognition of a pathogen-associated molecular pattern (PAMP; e.g. bacterial peptidoglycan), leading to the phosphorylation and degradation of the inhibitor of kappa B (IκB; encoded by *cactus* for Toll signalling, and by the *relish* C-terminus in Imd signalling) (reviewed in Valanne, Wang and Ramet, 2011; Myllymaki, Valanne and Ramet, 2014). Under non-signalling conditions, IκB sequesters NF-_k_B transcription factors in the cytoplasm. These transcription factors are encoded by *dorsal (dl)* and *Dorsal immune-related factor (Dif)* in Toll signalling, and *relish (rel)* in Imd signalling, and all translocate to the nucleus to induce gene expression following ΙκΒ degradation (reviewed in Valanne, Wang and Ramet, 2011; Myllymaki, Valanne and Ramet, 2014). Although the mechanism by which Toll and Imd recognise RNA viruses is unclear, both are active and provide immunity against some viral infections in insects, most likely through induction of antiviral effector responses. For example, Toll is broadly antiviral against RNA viruses such as Drosophila C virus, Nora virus, and Flock House virus in *Drosophila* during orally acquired, but not systemic infections, and in *Aedes* mosquitoes against dengue virus (Zambon *et al.*, 2005; Xi, Ramirez and Dimopoulos, 2008; Ramirez and Dimopoulos, 2010; Ferreira *et al.*, 2014). Additionally, Imd is antiviral against a subset of viruses in *Drosophila*, such as Cricket Paralysis virus, Drosophila C virus, and Sindbis virus and in *Aedes* cell culture against the alphaviruses Semliki Forest virus and O’nyong’nyong virus (Fragkoudis *et al.*, 2008; Avadhanula *et al.*, 2009; Costa *et al.*, 2009; Waldock, Olson and Christophides, 2012; Sansone *et al.*, 2015).

Although the effect of NF-_k_B signalling on DNA virus infection in insects has not been directly tested, polydnaviruses, ascoviruses, baculoviruses, and entomopoxviruses have acquired suppressors of NF-_k_B signalling by horizontal gene transfer, providing indirect evidence for anti-DNA virus NF-κB signalling (Thoetkiattikul, Beck and Strand, 2005; Lamiable *et al.*, 2016). First, a ‘polydnavirus’ encoded in the genome of the Braconid parasitoid wasp *Microplitis demolitor* has acquired homologs of ΙκΒ, some of which inhibit Dif and rel by direct binding (Thoetkiattikul, Beck and Strand, 2005). However, this is a domesticated endogenous viral element that forms a component of the wasp venom, and as these ΙκΒ homologues are not found in related nudiviruses, baculoviruses, or hytrosaviruses, it seems likely they were acquired to inhibit anti-parasitoid immune responses in the host of the parasitoid wasp, rather than the antiviral immune response of the wasp itself (Bitra, Suderman and Strand, 2012; Herniou *et al.*, 2013). Second, homologs of *diedel*, which encode a cytokine that inhibits the Imd pathway in *Drosophila*, are similarly found in ascoviruses, baculoviruses, entomopoxviruses, and *Leptopilina spp.* polydnavirus venom, likely through independent horizontal transfer from arthropod hosts (Lamiable *et al.*, 2016). Virus-encoded diedel phenocopies fly-encoded diedel, suggesting that viral diedel has retained an Imd-suppressive function, and that the Imd pathway likely interacts with these DNA viruses (Lamiable *et al.*, 2016). However, it is still unclear whether antiviral Toll signalling is targeted by insect virus-encoded immune suppressors, and whether these hijacked host pathway inhibitors represent a subset of a greater diversity of NF-_k_B immune inhibitors or reflect evasion of virus-specific immune mechanisms.

The recent isolation of Kallithea virus (KV; Webster *et al.*, 2015; Palmer *et al.*, 2018), a nudivirus that naturally infects *Drosophila melanogaster* at high prevalence in the wild, provides a tractable system to study host-DNA virus interactions and to identify immune evasion strategies in DNA viruses. Moreover, because some virus-encoded immune suppressors have been found to be highly host-specific, the use of native host-virus pairs is vital to our understanding of viral immune evasion (e.g. Parisien, Lau and Horvath, 2002; Mariani *et al.*, 2003; Goffinet *et al.*, 2009; Rajsbaum *et al.*, 2012; van Mierlo *et al.*, 2014; Stabell *et al.*, 2018). Here, we use this system to describe the interaction between antiviral immune pathways and a DNA virus in *Drosophila.* Using mutant fly lines, we find that the RNAi and Imd pathways mediate antiviral protection against KV *in vivo*, but that abrogation of Toll signalling has no effect on virus replication. Through re-analysis of previous RNA-sequencing data, we observe a broad downregulation of NF-_k_B responsive antimicrobial peptides following infection, and perform a small-scale screen for KV-encoded immune inhibitors. We identify viral protein gp83 as having a complex interaction with NF-_k_B signalling, leading to induction of Imd signalling but potent suppression of Toll signalling. This suppression acts directly through, or downstream of, NF-_k_B transcription factors. Finally, through deletions of conserved protein regions and analysis of the related *Drosophila innubila* nudivirus (DiNV) gp83 ortholog, we show that the immunosuppressive activity of gp83 against *D. melanogaster* NF-_k_B signalling is conserved.

## Materials and Methods

### Fly strains, virus growth, and mortality experiments

All fly lines were maintained and crossed on standard cornmeal medium at 25 °C. Viral titre and mortality were measured following KV infection in two control lines (*w*^1118^ and *Oregon R)* and in mutant lines compromised in the following immune signalling pathways: RNAi (*Dcr*-*2*^L811fsx^ (Lee *et al.*,2004) and *AGO2*^414^ Okamura *et al.*, 2004)), Toll (*spz*^4^ (Weber *et al.*, 2007), *dl*^*1*^ (Nusslein-Volhard, 1979), *Dif*^1^ (Rutschmann *et al.*, 2000), and p*ll*^2^/p*ll*^21^ trans-heterzygotes (Anderson and Nüsslein-Volhard, 1984; Hecht and Anderson, 1993)), and Imd (*rel*^e20^ (Hedengren *et al.*, 1999) and *imd*^*10191*^ (Pham *et al.*, 2007)).

For mortality assays, 100 female flies of each genotype were injected with 50 nL of either KV suspension (10^5^ ID_50_, as described in (Palmer *et al.*, 2018)) or chloroform-treated KV suspension (which inactivates KV through the destruction of the membrane) and transferred in groups of 10 to sucrose agar vials. The number of surviving flies was recorded on alternate days, and each group of flies was transferred to fresh food each week. Per-day mortality was analysed as a binomial response variable with the Bayesian generalised linear mixed modelling R package, MCMCglmm (Hadfield, 2010), with days post-inoculation (dpi), dpi^2^ (to allow for non-linear changes in mortality), and genotype as fixed effects, and vial as a random effect, as described previously (Palmer *et al.*, 2018). All confidence intervals are reported as 95% highest posterior density (HPD) intervals.

Viral titre was measured in each line after intra-abdominal injection of 50 nL of KV suspension. Infected female flies of each line (n=50) were transferred to 10 sucrose agar vials in groups of 5, and 5 vials of each genotype were homogenised in Trizol at 5 and 10 dpi. For RNAi mutants, flies were also assayed at 3 dpi. DNA was extracted by phenol-chloroform precipitation and viral titre estimated by quantitative PCR relative to host genomic DNA, using previously described primers (rpl32; Palmer *et al.*, 2018)). Log-transformed viral titre was analysed as a Gaussian response variable using MCMCglmm (Hadfield, 2010), with genotype, dpi, and genotype-by-dpi interactions as fixed effects. Titre in RNAi and NF-_k_B mutants were assayed in separate experiments, and therefore analysed independently. We took a statistical approach to account for the impact of differing genetic backgrounds between mutant lines, using the range of KV titres seen previously across 120 different natural genetic backgrounds from the *Drosophila* Genetic Reference Panel (Palmer *et al.*, 2018). Specifically, considering *w*^*1118*^ and *Oregon R* as controls and mutants of each pathway as the ‘experimental’ group, a null distribution of effect sizes expected only from differences in genetic background was created by randomly choosing two DGRP lines to serve as controls and additional DGRP lines reflecting the mutant lines used in each pathway. For each null draw, the same model was fitted as described above, the absolute value of the effect size was recorded, and this was repeated 1000 times to obtain a distribution. If the average effect size associated with mutants in a pathway was greater than the highest 5% of effect sizes, we concluded that the observed differences in KV titre were due to mutations in the tested pathway.

### Cell culture and virus propagation

S2 cells were cultured at 25 °C in Schneider’s Drosophila Medium with 10% heat-inactivated fetal bovine serum and 50 U/mL penicillin and 50 ug/mL streptomycin (Life technologies). KV was purified from flies 10 days after initial infection as previously described (Palmer *et al.*, 2018).

### Cloning

We selected 10 KV genes identified as highly expressed at three dpi (Palmer *et al.*, 2018) to screen for KV-encoded immune suppressors. These were *gp23, gp43, gp72, gp83, ACH96233.1-like, ACH96143.1-like, putative protein 1, putative protein 12, putative protein 15, putative serine protease* (corresponding to GenBank accession numbers AKH40365.1, AKH40394.1, AKH40346.1, AKH40369.1, AKH40392.1, AKH40340.1, AQN78560.1, AKH40392.1, AKH40404.1, and AQN78556.1). Each KV gene was amplified using the Qiagen Long Range PCR kit as per the manufacturer’s instructions, with primers that introduced restriction sites and the *Drosophila* Kozak sequence (restriction enzymes and primers used in Supplementary Table 1), and cloned into a pAc5.1 vector (Invitrogen) with a V5-His C-terminal tag. The KV gene *gp83* was also cloned into pAc5.1 vector with GFP instead of V5-His to introduce a C-terminal GFP tag. Deletion constructs for gp83 were created by separately amplifying 2 segments of gp83 with primers that span the desired deletion and performing a second PCR reaction with these segments as a template, and the forward and reverse primers from the 5’ and 3’ segments, respectively (Supplementary Table 1; gp83^Δ1^: CGLIECSELLRDRLCSKL deletion; gp83^Δ2^: WSDRLNLI deletion). The resulting amplicons with deletions were cloned as described above. The *gp83* gene from DiNV (Unckless, 2011; Hill and Unckless, 2017) was also cloned as above (Supplementary Table 1).

Additionally, Toll pathway components *pll, tube, cact, Dif*, and *dl* were cloned into the pAc5.1 vector, as described above (Supplementary Table 1). Other Toll and Imd pathway constructs have been described before, including pAc5.1-Toll^lrr^ (Tauszig *et al.*, 2000), pAc5.1-dl-GFP (Li and Dijkers, 2015), pMT-PGRP-LCx (Kaneko *et al.*, 2006), pAc5.1-rel-GFP (Foley and O’Farrell, 2004), and the firefly luciferase (FLuc) reporter plasmids with promoter sequences from *Drosomycin (Drs), Diptericin (Dpt)*, and *Attacin-A (Att-A)* (Tauszig *et al.*, 2000) or with 10X STAT binding sites (Baeg, Zhou and Perrimon, 2005).

### Transfection and RNAi Knockdown in S2 cells

S2 cells were transfected using Effectene transfection reagent, as per the manufacturer’s instructions. Double-stranded RNA (dsRNA) was synthesized against *cactus, gp83, FLuc, renilla luciferase (RLuc)*, and *GFP* to knockdown these genes in S2 cells. Primers with flanking T7 sequences were used to amplify regions of each gene (Supplementary Table 1) and dsRNA was synthesized from the resulting PCR products with T7 RNA polymerase and purified using GenElute Total RNA mini kit (Qiagen) (e.g. van Cleef *et al.*, 2011).

### Immune suppression assays

The 10 cloned KV genes were tested for their ability to suppress RNAi, JAK-STAT, Toll, or Imd activity. RNAi suppression assays were performed as described previously (van Cleef *et al.*, 2011). Briefly, 5×10^4^ S2 cells were seeded in a 96-well plate and 24 hours later transfected with 33 ng of pMT-FLuc, 33 ng pMT-Rluc, and 33 ng of either pAc5.1 empty vector or the pAc5.1 expression plasmid encoding a KV gene. Two days later, 400 ng of either GFP or GL3 dsRNA was added to each well, and CuSO4 was added 8 hours later to a final concentration of 500 μM to induce expression of the luciferase reporters. RLuc and FLuc luciferase activity were measured using the Dual Luciferase Assay Kit (Promega).

For JAK-STAT immunosuppression assays, 5×10^4^ S2 cells were seeded in a 96-well plate and transfected 24 hours later with 30 ng of 10XSTAT-FLuc, 20 ng pAc5.1-Rluc, and 50 ng of either pAc5.1 empty vector or the pAc5.1 expression plasmid encoding a KV gene. Luciferase activity was measured at 48 hours following transfection.

For NF-_k_B immunosuppression assays, a plasmid encoding the Imd receptor PGRP-LC (isoform x; pMT-PGRP-LCx) (Werner *et al.*, 2003; Kaneko *et al.*, 2006) or a constitutively active Toll construct lacking the extracellular leucine-rich repeat domain, pAc5.1-Toll^lrr^ (Tauszig *et al.*, 2000) was transfected alongside each KV gene, and a NF-_k_B-responsive FLuc reporter containing the either the *Dpt* (Imd) or *Drs* (Toll) promoter sequence (Tauszig *et al.*, 2000). For Toll immune suppression assays, 5×10^4^ S2 cells were seeded in 96-well plates and 24 hours later transfected with 50 ng of either empty pAc5.1 vector or a pAc5.1 KV gene expression construct, 20 ng of either pAc5.1 or pAc5.1-Toll^lrr^, 10 ng of Drs-FLuc, and 10 ng pAc5.1-Rluc. Imd immune suppression assays were performed in the same manner, except that pMT, pMT-PGRP-LCx, and Dpt-FLuc were substituted for pAc5.1, pAc5.1-Toll^lrr^, and Drs-FLuc, respectively, and CuSO4 was added immediately following transfection. Analogous experiments were performed using pAc5.1-dl, pAc5.1-Dif, and pAc5.1-pll instead of pAc5.1-Toll^lrr^, or by transfecting 5 ng of *cact* dsRNA. In the latter case, 70 ng of KV gene expression construct was transfected instead of 50 ng. RLuc and FLuc activity were assayed 48 hours after transfection.

Immunosuppression assays were also performed using KV-infected cells. 5×10^4^ cells were seeded in 96-well plates, followed by the immediate addition of 5 μL of either KV suspension (10^3^ ID50) or chloroform-treated KV, and transfected the next day. For RNAi suppression assays with KV, 50 ng pMT-RLuc, 50 ng pMT-FLuc (van Cleef *et al.*, 2011), and 5 ng of either GFP or GL3 dsRNA were transfected 2 dpi and CuSO_4_ added 8 hours later. To measure JAK-STAT activity following KV infection, 70 ng of 10XSTAT-FLuc and 30 ng pAc5.1-Rluc (Van Cleef *et al.*, 2014) were transfected. For Toll suppression assays, 70 ng of either pAc5.1 or pAc5.1-Toll^lrr^, 20 ng of Drs-FLuc, and 10 ng pAc-RLuc were transfected. Finally, to measure Imd activity following KV infection, 70 ng of either pMT or pMT-PGRP-LCx, 20 ng of Dpt-FLuc, and 10 ng pAc-RLuc were transfected, and CuSO4 was added immediately following transfection. Luciferase activity was measured at 4 dpi.

The R package MCMglmm was used to determine significance in immune suppression assays, with the RLuc-normalised FLuc values as a Gaussian response variable. In the original screen for immune suppressors, any experimental induction of an immune pathway was treated as a fixed effect (e.g. addition of dsRNA against FLuc in the RNAi suppression assay, PGRP-LC overexpression in the Imd suppression assay, and Toll^lrr^ in the Toll suppression assay), each KV gene was treated as a random effect, and the interaction between KV gene and the induced experimental change to signalling output was treated as a random effect. In subsequent NF-_k_B suppression experiments, where the only tested KV gene was gp83, gp83 and the interaction between gp83 and overexpression of NF-_k_B receptors were treated as fixed effects. Likewise, when immune suppression experiments were carried out with KV-infected cells instead of over-expressing KV genes, KV infection status, the induction of an immune pathway, and the interaction between these were treated as fixed effects.

### Immunoprecipitation and western blotting

To test whether gp83 directly interacted with dl, 2×10^6^ cells were seeded in 6-well plates and transfected with 150 ng of either pAc5.1 empty vector, pAc5.1 encoding V5-tagged gp83, or V5-tagged cact alongside 150 ng of the expression plasmid (pAc5.1) encoding GFP or GFP-tagged dl. Two days post-transfection, two wells per treatment were resuspended in lysis buffer (0.1% NP-40, 30 mM Hepes-KOH, 150 mM NaCl, 2mM MgOAc) supplemented with cOmplete protease inhibitor cocktail (Roche) and 5 mM DTT, and disrupted 30 times through a 25-gauge needle. After 10 minutes incubation on ice, cell debris was pelleted by centrifuging at 16,000xg for 30 minutes and supernatant was either stored as an input control or collected and incubated for 5 hours at 4 °C with magnetic control beads. Binding control beads were removed and the resulting supernatant was incubated with GFP-trap magnetic beads (Chromotek) overnight at 4 °C. Beads were washed 3 times in lysis buffer and 3 times in 25 mM Tris-HCl, 150 mM NaCl solution, and protein complexes eluted by boiling 10 minutes at 95 °C in Laemmli buffer.

Whole cellular protein extracts were prepared by heating S2 cells for 10 min at 95 °C in Laemmli buffer. Whole cellular extracts or immunoprecipitated proteins were separated on a 12% SDS-PAGE gel and transferred to a nitrocellulose membrane. Non-specific binding was blocked with blocking solution (phosphate buffered saline with 0.1% Triton-X (PBT) and 5% dry milk). Proteins of interest were probed with primary antibody diluted in blocking solution overnight at 4 °C, and visualized with an hour incubation of secondary antibody in blocking solution. Membranes were washed 3 times in PBT before and after each step. Antibodies that were used include mouse anti-dl (1:100 dilution, Developmental Studies Hybridoma Bank), mouse anti-V5 (1:1000 dilution, Invitrogen), rat anti-tub-α (1:1000 dilution, SanBio), and rabbit anti-GFP (1:1500 dilution, abcam ab6556) as primary antibodies, and goat anti-mouse IR-Dye 680 (1:15,000 dilution, LI-COR), goat anti-rat IR-Dye 800 (1:15,000 dilution, LI-COR), goat anti-rabbit IR-Dye 800 (1:15,000, LI-COR). An Odyssey Infrared Imager (LI-COR) was used to image blots.

### Mass spectrometry

S2 cells were seeded (10^6^) and co-transfected with pCoBLAST and pAc5.1-gp83^gfp^ plasmid at a 1:19 ratio (125 ng and 2.38 μg, respectively). Medium was replaced 3 hours post-transfection, and again at 48 hours post-transfection with medium supplemented with blasticidin (20 μg/mL). Another 48 hours later, cells were refreshed with medium containing 4 μg/mL blasticidin, which was thereafter replaced every 3-4 days with medium containing 4 μg/mL blasticidin, resulting in a polyclonal cell line.

For mass spectrometry, wild-type S2 cells or S2 cells stably expressing GP83^gfp^ were lysed in 50mM Tris-HCl (pH 7.8), 150mM NaCl, 1% NP-40, 0.5mM DTT, 10% glycerol and protease inhibitor cocktail (Roche). Approximately 4 mg of protein lysate was subjected to GFP-affinity purification using 7.5 μL GFP-trap beads (Chromotek) for approximately 1.5 hours at 4 °C. Beads were washed twice in lysis buffer, twice in PBS containing 1% NP-40, and three times in PBS, followed by on-bead trypsin digestion as described previously (Smits *et al.*, 2013). Afterwards, tryptic peptides were acidified and desalted using Stagetips, eluted, and brought onto an EASY-nLC 1000 Liquid Chromatograph (Thermo Scientific). Mass spectra were recorded on a QExactive mass spectrometer (Thermo Scientific) and MS and MS2 data were recorded using TOP10 data-dependent acquisition. Maxquant (v1.5.1.0) was used to analyse raw data, using recommended settings (Cox and Mann, 2008). LFQ, IBAQ, and match between runs were enabled. The peptides were mapped to *Drosophila melanogaster* proteins (UniProt June 2017) and contaminants and reverse hits were filtered with Perseus (v1.3.0.4) (Tyanova *et al.*, 2016). Missing values were imputed, assuming a normal distribution, and significance determined by a t-test on log-transformed LFQ-values between wild-type and gp83-expressing S2 cells.

### Immunofluorescence microscopy

5×10^5^ S2 cells were seeded in 12-well plates with glass coverslips in each well. Cells were transfected with 100 ng of pAc5.1 or pAc5.1-gp83-V5 and 100 ng of pAc5.1-dl-GFP. Two days after transfection, cells were fixed with 4% paraformaldehyde, washed twice in PBS, once with PBT, and blocked with PBT with 10% goat serum. Cells were stained by incubation with mouse anti-V5 (1:400, Invitrogen) for one hour, followed by fluorophore-containing goat anti-mouse secondary antibody (1:400, AlexaFluor) with 10 ug/mL Hoechst for one hour. Finally, cells were washed twice in PBT and twice in PBS, mounted on slides with Fluoromount-G (eBiosciences), and imaged with an Olympus FluoView FV1000. Fluorescence was measured in whole cells, or separately in the cytoplasm and nuclei by outlining the region of interest in Fiji (Schindelin *et al.*, 2012) to calculate the mean fluorescence.

## Results and Discussion

### RNAi and Imd pathways are antiviral against KV in vivo

The RNAi pathway provides antiviral activity against the DNA virus IIV6, and KV-derived vsiRNAs are produced upon infection of adult naturally-infected *Drosophila* (Bronkhorst *et al.*, 2012, 2014; Webster *et al.*, 2015). However, the contribution of Imd and Toll pathways to anti-DNA virus immunity have not been described. We used fly lines mutant for RNAi, Imd, and Toll pathway components to assess whether these pathways fulfil an antiviral function during KV infection. First, we infected mutants for RNAi genes *Dcr-2* and *AGO2* with KV, and measured viral titre and mortality following infection. Following KV infection, both *Dcr-2* and *AGO2* mutants exhibited significantly greater KV titres at 3 dpi, with KV titre 78-fold greater in *Dcr-2* mutants (95% HPD intervals: 18-281 fold; MCMCp < 0.001) and 55-fold greater in *AGO2* mutants (13-237 fold, MCMCp < 0.001; Figure 1). However, the increased KV replication in RNAi mutants was not sustained at later infection timepoints. At 5 dpi, *Dcr-2* mutants did not have significantly different KV titre from the controls (MCMCp = 0.22), and KV had a slightly diminished advantage in *AGO2* mutants (12-fold increase; 2.5-43 fold, MCMCp < 0.001; Figure 1). By 10 dpi, there was no significant difference between viral titre in control flies and either *Dcr-2* mutants (MCMCp = 0.43) or *AGO2* mutants (MCMCp = 0.7). Therefore, either the antiviral effect of RNAi is short-lived (for example, a viral suppressor of RNAi may eventually be expressed *in vivo)*, other immune pathways take over as the dominant antiviral force, or KV negatively regulates its own replication or saturates a resource. Nevertheless, despite the similar titres during late infection, there was still a significant increase in KV-induced mortality in *Dcr-2* and *AGO2* mutants, where 70% of control flies were alive at 19 dpi, compared to 25% in *Dcr-2* mutants (MCMCp < 0.001) and 38% in *AGO2* mutants (Figure 1), possibly due to early host damage or increased expression of virulence factors throughout infection (e.g. Jayachandran, Hussain and Asgari, 2012). These results extend the antiviral role of the RNAi pathway to KV infection.

**Figure 1:**
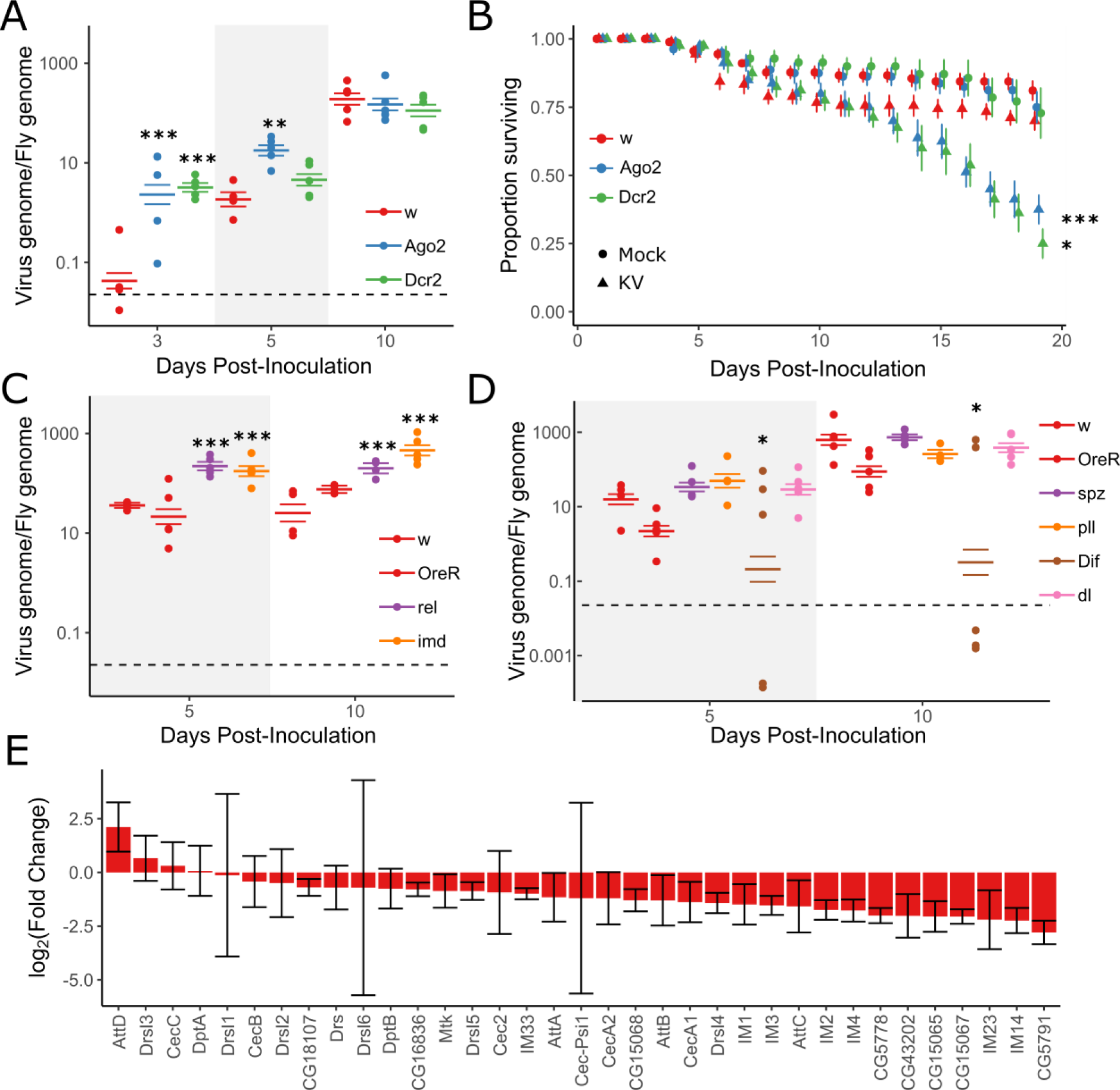
RNAi and Imd pathways provide antiviral defense against Kallithea virus. Mutants for RNAi (A,B) and NF-_k_B (C,D) pathways were assayed for viral titre (A,C,D) and mortality (B) following KV infection. Viral titre was measured by qPCR, relative to rpl32 DNA, where each point is a vial of 5 flies, and coloured horizontal lines correspond to the mean titre and associated standard error (A,C,D). Horizontal dotted lines (A,C,D) represent the amount of virus injected at the zero timepoint. (A) RNAi mutants have increased early viral titre relative to control *w*^1118^ flies at 3 dpi, but no difference in titre by 10 dpi. (B) RNAi mutants exhibit increased mortality following KV infection, relative to chloroform-treated KV controls (mock). Each point is the mean number of surviving flies across 10 vials of 10 flies, with associated standard errors. (C) *Imd* and *rel* mutant flies have increased viral titre relative to two wild-type lines (*w*^1118^ and OreR) at 5 and 10 dpi. (D) However, Toll pathway mutants for *spz, pll, Dif*, or *dl* show no consistent difference in KV titre relative to control lines at 5 or 10 dpi. (E) Genes encoding Toll and Imd-responsive antimicrobial peptides are generally downregulated following KV infection of *OreR* flies 3 dpi, relative to uninfected controls (RNA-sequencing data from Palmer et al, 2018; ERP023609; n = 5 libraries per treatment), consistent with the presence of a virus-encoded immune inhibitor. Error bars show standard errors of the mean. *p < 0.05; **p < 0.01; ***p < 0.001. (Statistical tests performed in MCMCglmm)

We next infected Imd and Toll pathway mutants with KV and assessed KV DNA levels by qPCR at 5 and 10 dpi. We found Imd pathway mutants had significantly greater viral titre as compared to two control lines, with *imd* mutants having 6-fold greater KV titre at 5 and 10 dpi (2.7-13.7 fold, MCMCp<0.001), and *rel* having 8-fold greater viral titre at 5 and 10 dpi (3.1-15.9 fold, MCMCp < 0.001; Figure 1). Because the Imd effect spans 5 and 10 dpi, and we have previously measured KV titre in 125 inbred lines of the *Drosophila* Genetic Reference Panel at 8 dpi (Mackay *et al.*, 2012; Palmer *et al.*, 2018), we attempted to account for genetic background by comparing the average effect of Imd mutants to the distribution of effects consistent with natural variation in the genetic background. This analysis indicated that the increased titre observed in Imd mutants is unlikely to be due to genetic background (p = 0.01), if the variation in the DGRP is representative of the variation expected between lab-maintained fly lines. We also infected flies mutant for the Toll pathway components *spz, pll, Dif*, and *dl.* Viral titre was unchanged in most Toll pathway mutants compared to controls, except in *Dif* mutants (MCMCp = 0.02; Figure 1), where some vials remained uninfected. Further, the pathway-level effect of Toll mutants was within the expected distribution of effects caused by differences in genetic background, even when *Dif* mutants were excluded (p = 0.28). We concluded that the Imd pathway is antiviral against KV, but that abrogation of Toll function has no effect on KV growth. This could indicate that Toll is not antiviral against this DNA virus, or that it is efficiently suppressed by virus infection. The latter is consistent with our observation that genes encoding antimicrobial peptides are generally downregulated in KV-infected flies compared to uninfected controls (Figure 1), and we therefore explored the capability of KV to suppress innate immune pathways in cell culture.

### KV growth in cell culture

KV growth in *D. melanogaster* cell culture has not previously been described. We found variation in the ability of KV to infect five commonly-used cell lines, and KV grew well in S2, S2R+, and DL2 cells, but poorly in Kc167 and Dm-BG3-c2 cells (Figure S1). In S2 cells, which we used for further analyses, KV was released into the medium at substantial levels starting from 3 dpi (Figure S1). Therefore, in all subsequent experiments, we assayed cells at 4 dpi, assuming that a high proportion of cells would be infected at this timepoint. We did not observe any overt cytopathic effects of KV-infected cells within 14 days of infection. However, when KV-infected cells were passaged at 7 dpi, we observed larger (MCMCp < 0.001) and fewer (MCMCp < 0.001) cells, likely due to a decrease in cell proliferation (Figure S1).

### KV inhibits JAK-STAT and Toll, and induces Imd signalling in cell culture

We used previously established luciferase reporter-based assays to describe the effect of KV infection on RNAi, JAK-STAT, Toll, and Imd pathways. To determine if KV suppresses RNAi, we measured the RNAi silencing efficiency of cells inoculated with KV or chloroform-inactivated KV (hereafter referred to as mock-treated) by co-transfecting expression plasmids encoding FLuc with either GFP dsRNA or FLuc dsRNA. In both mock and KV-treated cells, FLuc dsRNA caused a 95% reduction in FLuc activity compared with GFP dsRNA treated cells, indicating KV infection does not inhibit RNAi in cell culture (MCMCp = 0.9; Figure 2). Many viruses studied in *Drosophila* encode a suppressor of RNAi (e.g. Li, Li and Ding, 2002; Van Rij *et al.*, 2006; Nayak *et al.*, 2010; van Mierlo *et al.*, 2012, 2014; Bronkhorst *et al.*, 2014; Van Cleef *et al.*, 2014), and therefore the absence of KV-induced RNAi suppression is somewhat surprising. It is possible that KV-RNAi interactions are different in the cell types that are naturally infected by KV, and that our inability to observe RNAi suppressive activity is a limitation of the cell culture model. Alternatively, if KV transmission does not occur until later stages of infection, there may be limited selective pressure to evade RNAi, as RNAi mutants and control flies have similar titres during late infection.

**Figure 2:**
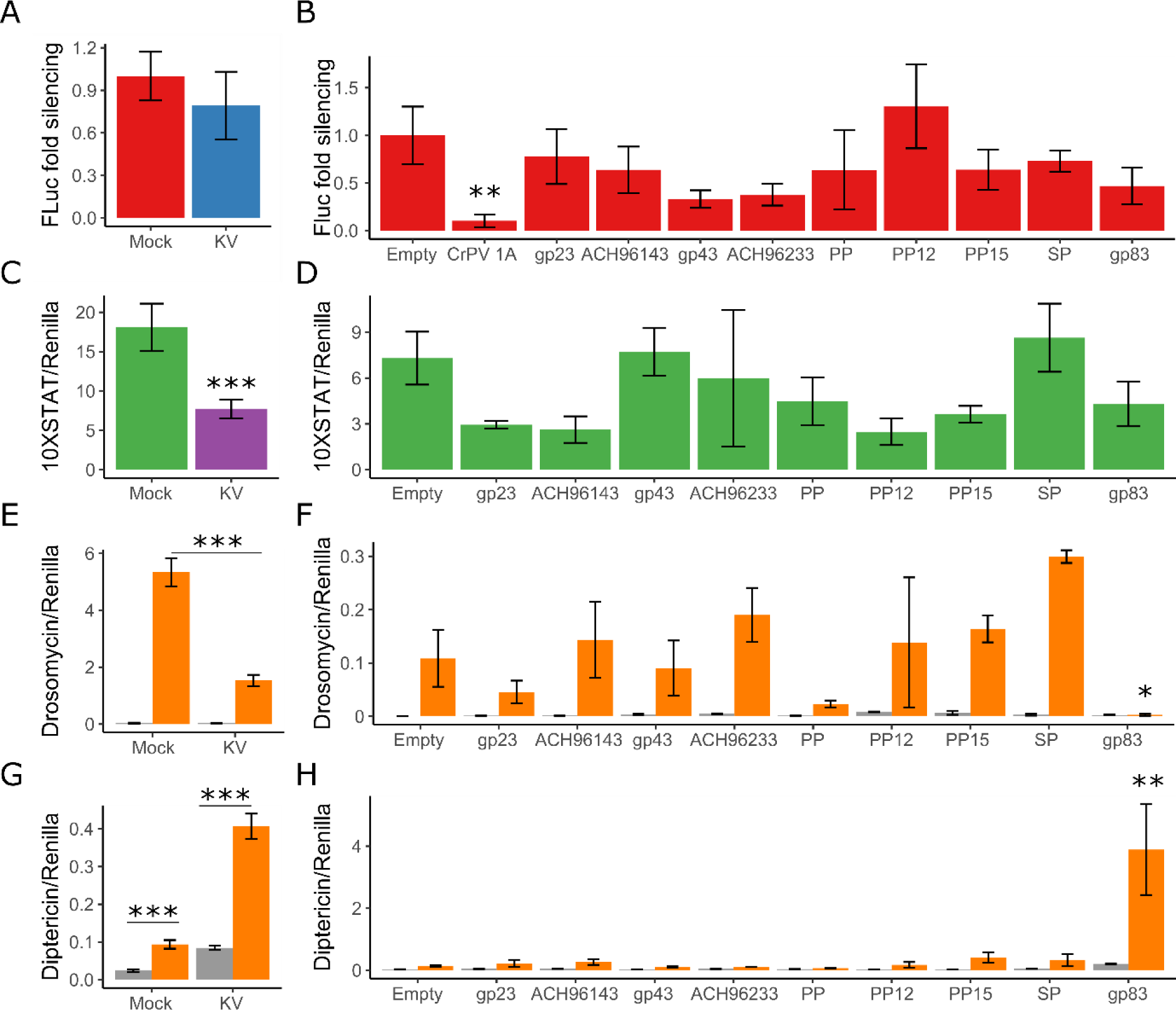
Kallithea virus gp83 suppresses Toll and induces Imd signalling. We assessed the ability of KV to inhibit RNAi, JAK-STAT, Toll, and Imd pathways 4 dpi (A,C,E,G), and whether 10 highly expressed KV genes interacted with these pathways (B,D,F,H). For RNAi suppression assays (A,B), fold silencing is the relative FLuc activity in cells treated with GFP dsRNA relative to those treated with FLuc dsRNA, normalised to 1 in mock-infected cells. For JAK-STAT suppression assays (C,D), cells were transfected with 10XSTAT-FLuc to measure JAK-STAT activation. For Toll suppression assays (E,F), cells were transfected with the Drs-FLuc reporter, with either pAc5.1 (Empty) or pAc5.1-Toll^lrr^. For Imd suppression assays (G,H), cells were transfected with the Dpt-FLuc reporter, with either pMT (Empty) or pMT-PGRP-LC. All luciferase values are relative to the constitutively expressed RLuc. (A) KV was unable to effectively inhibit RNAi silencing, as incubation with FLuc dsRNA was able to efficiently silence FLuc during KV infection. (B) Additionally, no tested KV gene inhibited RNAi (data combined from 2 experiments). (C) KV infection reduced JAK-STAT signalling in S2 cells. (D) However, overexpression of any of the 10 highly expressed KV genes was not significantly associated with STAT suppression. (E) Toll is not endogenously active in S2 cells (grey bars) but overexpression of Toll^lrr^ resulted in a dramatic increase in Drs luciferase (orange bars), which KV partially inhibited. (F) Overexpression of gp83 was able to completely inhibit Toll-induced Drs expression. (G) Likewise, Imd is not active in S2 cells, and overexpression of PGRP-LC led to increased Dpt luciferase (compare grey and orange bars). KV infection significantly induced Dpt luciferase with or without PGRP-LC. (H) Additionally, gp83 overexpression potently induced Imd signalling when coupled with PGRP-LC overexpression. PP=Putative Protein; SP=Serine Protease. Error bars show standard errors of the mean, calculated from 5 biological replicates for (A,C,E,G) and 3 biological replicates for (B,D,F,H). *p < 0.05; **p < 0.01; ***p < 0.001 (Statistical tests performed in MCMCglmm)

The JAK-STAT pathway has an antiviral role during Drosophila C virus infection (Dostert *et al.*, 2005) and mediates tolerance to the DNA virus IIV6, evidenced by upregulation of *vir-1* and the *Turandot (Tot)* genes (West and Silverman, 2018). However, previous transcriptional profiling did not identify differential expression of these genes following infection with KV (Palmer *et al.*, 2018). We assessed JAK-STAT activity in mock and KV-treated cells with a FLuc reporter containing ten STAT binding sites (Baeg, Zhou and Perrimon, 2005). This reporter is endogenously active in S2 cells (Baeg, Zhou and Perrimon, 2005), but KV infection led to a 58% reduction in STAT-mediated FLuc activity (37-74%, MCMCp < 0.001; Figure 2), indicating that JAK-STAT is down-regulated or inhibited following KV infection. However, in addition to mediating a transcriptional immune response, the JAK-STAT pathway is involved in cell proliferation (Arbouzova and Zeidler, 2006), which also decreases following KV infection in cell culture (Figure S1), making cause and effect difficult to distinguish.

We also assayed the effect of KV on Toll and Imd signalling. However, these pathways are not constitutively active in S2 cells. To measure KV suppression of these pathways, we therefore co-transfected Toll^lrr^ (a Toll receptor lacking the leucine-rich repeat extracellular domain) or PGRP-LC (an Imd pathway receptor) with *Drs* or *Dpt* luciferase reporters to artificially induce signalling of Toll and Imd pathways, respectively. Transfection of Toll^lrr^ increased Drs-Fluc by 243-fold (MCMCp < 0.001), consistent with previous reports (Tauszig *et al.*, 2000). However, KV infection reduced the maximum level of Toll^lrr^-mediated Drs activity by 81% (38-93%, MCMCp < 0.001; Figure 2), indicating KV can inhibit Toll signalling. Over-expression of PGRP-LC led to a 4-fold increase in Dpt-FLuc (3-5 fold, MCMCp < 0.001). In contrast to the effect on Toll signalling, KV infection led to a 3.6-fold increase (2.6-4.8 fold, MCMCp < 0.001) in Dpt-FLuc, which additively increased Dpt-FLuc when PGRP-LC overexpressing cells were infected with KV (17-fold increase compared to Imd-inactive, mock-treated cells; 12-23 fold, Figure 2). This suggests that KV infection in S2 cell culture suppresses Toll signalling but induces Imd signalling.

### KV-encoded gp83 modifies NF-_k_B signalling during infection

The immunosuppressive function of nudivirus genes has not previously been explored. To identify KV-encoded immune inhibitors, we cloned 10 KV genes that are highly expressed at 3 dpi in adult flies (Palmer *et al.*, 2018) and performed immune suppression assays for RNAi, JAK-STAT, Toll, and Imd pathways. We were unable to identify KV-encoded suppressors of RNAi or JAK-STAT among these 10 genes, although we confirmed that Cricket Paralysis Virus protein 1A potently suppressed RNAi in these assays (Nayak *et al.*, 2010); MCMCp = 0.006; Figure 2). However, we found that gp83— a KV gene encoding no recognisable protein domains—significantly reduced Toll^lrr^-induced Drs-FLuc expression (Figure 2). In this experiment, Toll^lrr^ overexpression induced Drs-FLuc by 24-fold (8-66 fold), but by only 1.9-fold (0.3-8 fold; MCMCp = 0.02) when gp83 was co-expressed. We confirmed that this was not an artefact of the luciferase reporter-based assay by repeating the experiment with qPCR of endogenous *Drs* as a readout, where gp83 overexpression potently reduced Toll^lrr^-induced *Drs* expression (MCMCp < 0.001; Figure S2). We further found that overexpression of gp83 caused a 5-fold (1.5-18 fold) *increase* in Imd-mediated Dpt-FLuc expression, with or without PGRP-LC overexpression (MCMCp = 0.008; Figure 2).

We next aimed to confirm that the interactions between the transfected KV gene gp83 and NF-_k_B pathways are representative of the function of gp83 during KV infection. Therefore, we silenced gp83 during KV infection using dsRNA, and measured associated changes in Toll, Imd, and JAK-STAT signalling. Co-transfection of gp83 with independent dsRNAs targeting gp83 completely reversed inhibition of Drs-FLuc compared with transfection of GFP dsRNA, indicating that these dsRNAs effectively silence gp83 (MCMCp < 0.001 for both dsRNAs; Figure S3). KV infection had no effect on Drs-FLuc (MCMCp = 0.26), but inhibited Toll^lrr^-induced signalling (MCMCp < 0.001), as reported above (Figure 2). However, knockdown of gp83 during KV infection led to increased Drs-FLuc in Toll-inactive cells (MCMCp = 0.004) and cells expressing Toll^lrr^ (MCMCp < 0.001; Figure 3). Likewise, knockdown of gp83 in KV-infected cells caused a modest decrease in Dpt-FLuc in Imd-inactive cells (MCMCp = 0.03), and this effect was much stronger in PGRP-LC overexpressing cells (MCMCp = 0.006; Figure 3). Consistent with a specific interaction with NF-_k_B signalling, gp83 knockdown had no effect on the ability of KV to suppress JAK-STAT signalling (MCMCp = 0.63; Figure 3).

**Figure 3:**
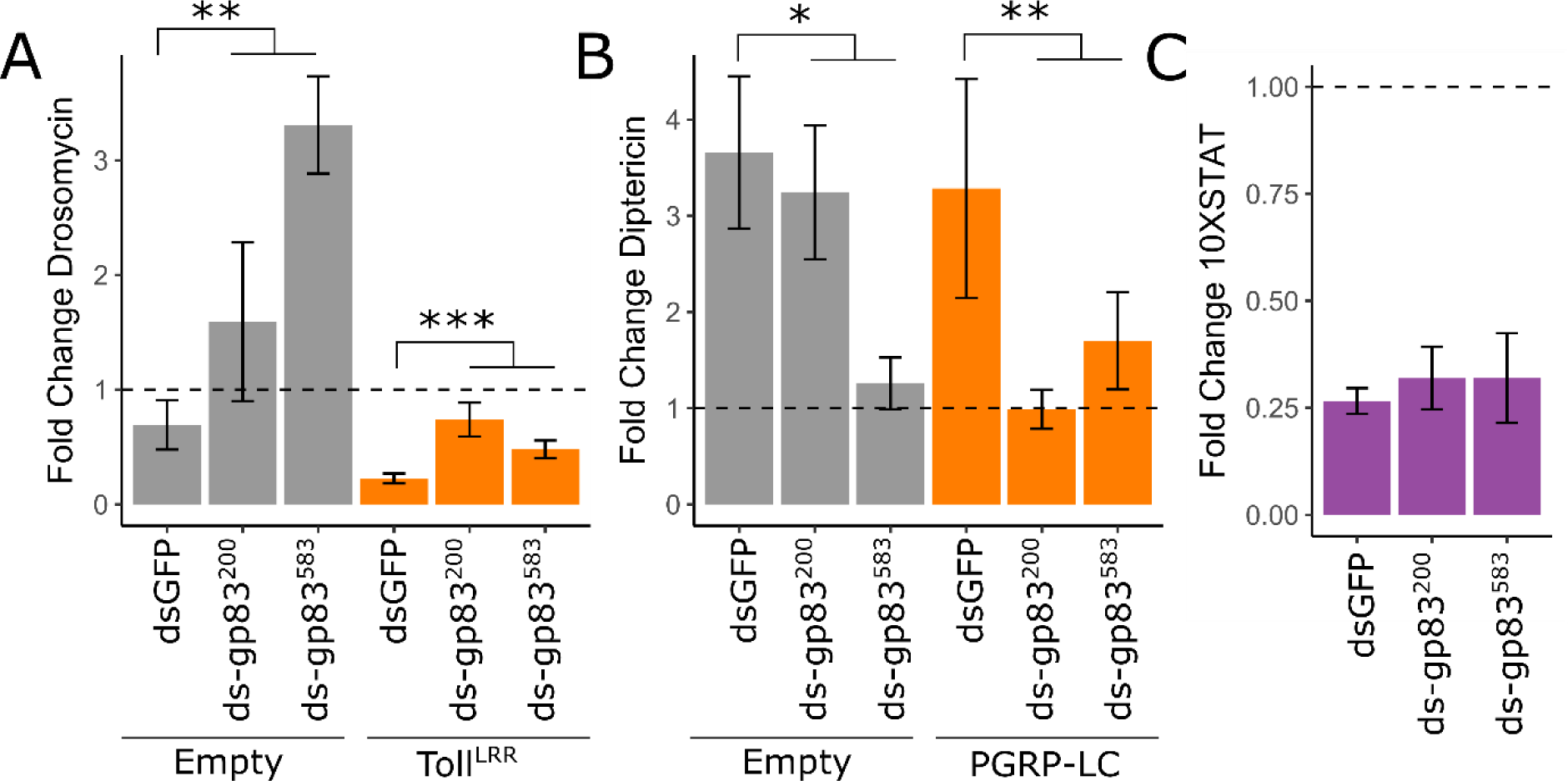
KV induction and suppression of NF-KB pathways is mediated by gp83. We assayed the ability of KV to inhibit Toll (A), induce Imd (B), and inhibit JAK-STAT (C) during gp83 knock down, using two independent dsRNAs against gp83 (labelled gp83 ^200^ and gp83^583^; see Figure S4 for confirmation of gp83 knockdown efficiency). Fold change in drosomycin, diptericin, and 10X-STAT was measured as Drs-FLuc, Dpt-FLuc, and 10XSTAT-FLuc activity, relative to RLuc expression. For each, fold-change in signalling following KV infection (4 dpi) is plotted, where 1 (horizontal dotted line) represents no induction or suppression of the pathway relative to mock infection (chloroform treated KV). (A) Knock-down of gp83 caused a significant increase in Drs luciferase expression following KV infection (grey bars) and significantly reduced the extent of KV suppression of Toll signalling during Toll^lrr^ overexpression (orange bars). (B) Knock down of gp83 causes significantly decreased Dpt luciferase expression following KV infection, especially during increased Imd activation (PGRP-LC overexpression; orange bars). (C) Knockdown of gp83 had no effect on the reduced JAK-STAT activity during KV infection (Figure 2), indicating gp83 interacts specifically with NF-_k_B signalling during KV infection. Error bars show standard error of the mean (n = 5 biological replicates). *p < 0.05; **p < 0.01; ***p < 0.001 (Statistical tests performed in MCMCglmm)

The immunosuppressive function of gp83 on Toll signalling is consistent with the observed downregulation of AMPs following KV infection *in vivo* and substantiates the hypothesis that Toll is antiviral and suppressed during infection. However, the induction of antiviral Imd signalling by a single viral protein is unexpected, and it is unclear why KV might not have evolved to avoid or supress Imd activation as seen for in other insect-infecting DNA viruses (Lamiable *et al.*, 2016). Assuming Imd activation is detrimental to virus transmission, this could indicate a trade-off between suppression of Toll and activation of Imd, or that gp83 is directly recognised by the fly immune system. We conclude that KV-encoded gp83 is involved in mediating complex interactions with NF-_k_B signalling, including suppression of Toll signalling and induction of Imd signalling.

### Immune suppression by gp83 occurs downstream of Toll transcription factors

Previously described polydnavirus-encoded Toll pathway inhibitors imitate ΙκΒ, blocking the nuclear entry of NF-_k_B transcription factors (Thoetkiattikul et al, 2005). Although the precise mechanism of interaction between gp83 and Toll signalling is unknown, suppression of Toll^lrr^-induced signalling indicates that gp83 also interferes with intracellular Toll signalling. We therefore performed genetic interaction experiments to narrow down the point in the Toll signalling pathway at which gp83 acts. Drs-FLuc was greatly increased by overexpressing *pll* (240-fold [131-414] induction of Drs-FLuc), silencing *cact* (75-fold [33-161] induction of Drs-FLuc), and overexpressing *Dif* (563-fold [317-1002] induction of Drs-FLuc;) or *dl* (459-fold [257-778] induction of Drs-FLuc;). Co-overexpression of gp83 potently reduced Drs-FLuc in each of these scenarios (MCMCp < 0.001 for each) - pll/gp83 co-overexpression led to a 0.55-fold change in Drs-FLuc (0.31-0.99 fold), cact^dsRNA^/gp83 led to a 1.73-fold change in Drs-FLuc (0.75-3.5 fold), Dif/gp83 led to a 0.86-fold change in Drs-FLuc (0.5-1.5 fold), and dl/gp83 led to a 1.5-fold change in Drs-FLuc (0.9-2.5 fold) relative to baseline Drs-FLuc expression (Figure 4). Additionally, V5 staining of gp83^V5^ revealed that gp83 is a nuclear protein (Figure 4E). Together, these results indicate that gp83 either inhibits NF-_k_B transcription factors, or acts downstream of them.

**Figure 4:**
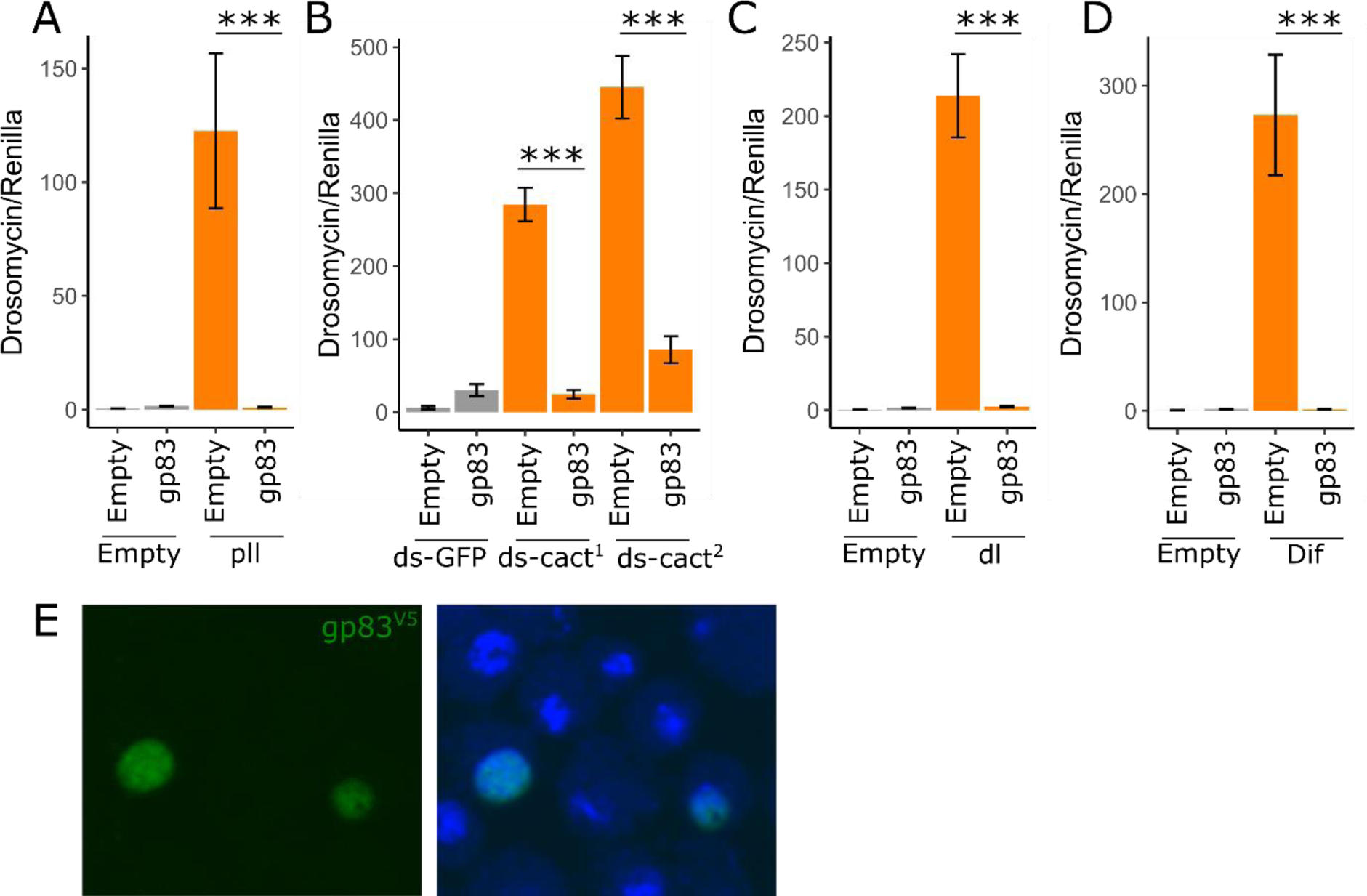
gp83 inhibits Toll signalling downstream of Dif and dorsal. Overexpression of *pll* (A), knockdown of *cactus* with two independent, non-overlapping dsRNAs (B), and overexpression of NF-_k_B transcription factors *dl* and *Dif* (C,D) led to increased Drs-FLuc expression, relative to RLuc expression, in S2 cells. In each case, gp83 was able to inhibit signalling, indicating that gp83 inhibits Toll signalling at the level of, or downstream of the NF-_k_B transcription factors. (E) Representative confocal image of S2 cells expressing V5-tagged gp83 stained with a V5 antibody (left pane) and a merged image in which nuclei are stained with Hoechst (right panel). Consistent with a downstream function, gp83 is a nuclear protein. Error bars show standard error of the mean (n = 5 biological replicates). *p < 0.05; **p < 0.01; ***p < 0.001 (Statistical tests performed in MCMCglmm)

Virus-encoded inhibitors of NF-_k_B in mammals have been reported to operate by promoting degradation of NF-_k_B transcription factors, blocking NF-_k_B access to the nucleus, or interfering with transcriptional co-activators to evade the interferon response (reviewed in Zhao *et al.*, 2015). Using co-immunoprecipitation and subsequent western blotting or mass spectrometry, we tested whether gp83 directly binds dl. Following immunoprecipitation of GFP-tagged dl, we were able to detect cact as an interacting positive control, but not gp83 (Figure S4). Additionally, we created an S2 cell line stably expressing GFP-tagged gp83, immunoprecipitated gp83^gfp^, and performed mass spectrometry on interacting partners. We identified 19 *Drosophila melanogaster* proteins, including 4 nuclear proteins (Nipped-B, Brf, Mlf, Ulp1), that were enriched in the gp83 immunoprecipitate (log2 fold enrichment > 2.5; FDR < 0.1; Figure S4). While we did not identify known downstream NF-_k_B pathway factors, the extracellular Toll ligand spz was among those enriched, despite the nuclear localization of gp83. However, dsRNA knockdown of spz did not rescue the immunosuppressive effect of gp83, indicating this interaction may not occur in live cells, or that it is not required for gp83 to inhibit Toll signalling. Further, knockdown of a subset of the enriched genes, including 3 of the 4 identified nuclear proteins, was unable to rescue the gp83 immunosuppressive effect (Figure S4), suggesting that gp83 may not form stable complexes with host proteins to interfere with NF-_k_B signalling (Figure S4).

Although we did not detect a direct association between dl and gp83, we observed a reduction in dl protein levels upon gp83 overexpression that is not dependent on Toll signalling (Figure S5). We quantified this effect by transfecting either GFP or GFP-tagged dl, with or without gp83, and measuring fluorescence by confocal microscopy. We found that while gp83 caused a 53% reduction in GFP levels, possibly due to a dl binding site in the actin 5C promoter of this construct (Zehavi *et al.*, 2014); 42-62%, MCMCp < 0.001), it caused a significantly greater reduction in dl^gfp^ (73% reduction; 66-78%, MCMCp < 0.001). However, KV infection did not decrease dl protein levels, indicating this may not be the primary mechanism by which KV inhibits Toll signalling (Figure S5). Instead, we hypothesize that gp83 interferes with the access of dl to either the nucleus or NF-_k_B binding sites, which indirectly affects dl localization and results in increased turnover. We prefer the latter explanation, that gp83 directly interferes with the Toll pathway transcriptional response, because overexpression of gp83 simultaneously induced the *Dpt* reporter (Figure 2) and reduced dl-responsive promoters (Drs-FLuc and Act5C-GFP; Figure 2, Figure S5). These observations implicate gp83 in regulating transcription at diverse loci responsive to both dl and rel, and suggest an interaction between gp83 and NF-_k_B-responsive genes.

### Immunosuppressive function of gp83 depends on conserved residues and is conserved in other nudiviruses

Conflict between the host immune system and virus-encoded immune inhibitors may result in an evolutionary arms race, leading to recurrent positive selection and eventual host specialization (e.g. Obbard *et al.*, 2009; Sawyer and Elde, 2012; Brockhurst *et al.*, 2014). Consistent with this, some immune inhibitors are only effective against their native host species, thereby defining the viral host range (e.g. Parisien, Lau and Horvath, 2002; Mariani *et al.*, 2003; Goffinet *et al.*, 2009; Rajsbaum *et al.*, 2012; van Mierlo *et al.*, 2014; Stabell *et al.*, 2018). We tested whether the immunosuppressive function of gp83 is conserved, and whether gp83 acts in a species-specific manner. The gp83 locus is absent from nudiviruses distantly related to KV, such as Heliothis zea nudivirus 1 (HzNV1), Tipula oleracea nudivirus (ToNV) and Peneaus monodon nudivirus (PmNV), but gp83 homologs are found in the more closely related Gryllus bimaculatus nudivirus (GrBNV), Nilaparvata lugens endogenous nudivirus (NlENV), Oryctes rhinoceros nudivirus (OrNV), Drosophila innubila nudivirus (DiNV), Tomelloso virus (TV), Mauternbach virus (MV), and Esparto virus (EV; Figure 5). Although gp83 lacks recognisable protein domains, several regions are strongly conserved among these nudiviruses, suggesting functional conservation (Figure 5). To test whether gp83 function depends on these conserved domains, we made two gp83 deletion constructs (gp83^Δ1^ and gp83 ^Δ2^) that respectively remove conserved regions of 18 and 8 amino acids without substantially altering protein stability, and transfected these alongside Toll^lrr^ with the Drs-FLuc reporter. Although detectable by western blotting, gp83 ^Δ1^ (MCMCp = 0.67) or gp83 ^Δ2^ (MCMCp = 0.79) were unable to inhibit Toll signalling, indicating that these conserved residues are important for the immunosuppressive function of gp83. To test whether gp83 function is conserved among viruses, we cloned gp83 from DiNV and performed Toll immunosuppression assays. The gp83 homolog from DiNV was able to completely inhibit *D. melanogaster* Toll signalling (MCMCp < 0.001), despite only 57% amino acid identity with KV gp83, demonstrating that the immunosuppressive function of gp83 is conserved in other nudiviruses and that it is not highly host-specific. This observation suggests that the Toll-gp83 interaction may not be a hotspot of antagonistic ‘arms race’ coevolution and has not led to specialization of DiNV gp83 to the *D. innubila* immune system at the expense of its ability to function in *D. melanogaster.* This could be because gp83 has relatively few direct interactions with host proteins (Figure S4), and may instead interact directly with transcription factor binding sites which are under high constraint, and therefore unable to evolve resistance to the immunosuppressive effect of gp83 (Nitta *et al.*, 2015).

**Figure 5:**
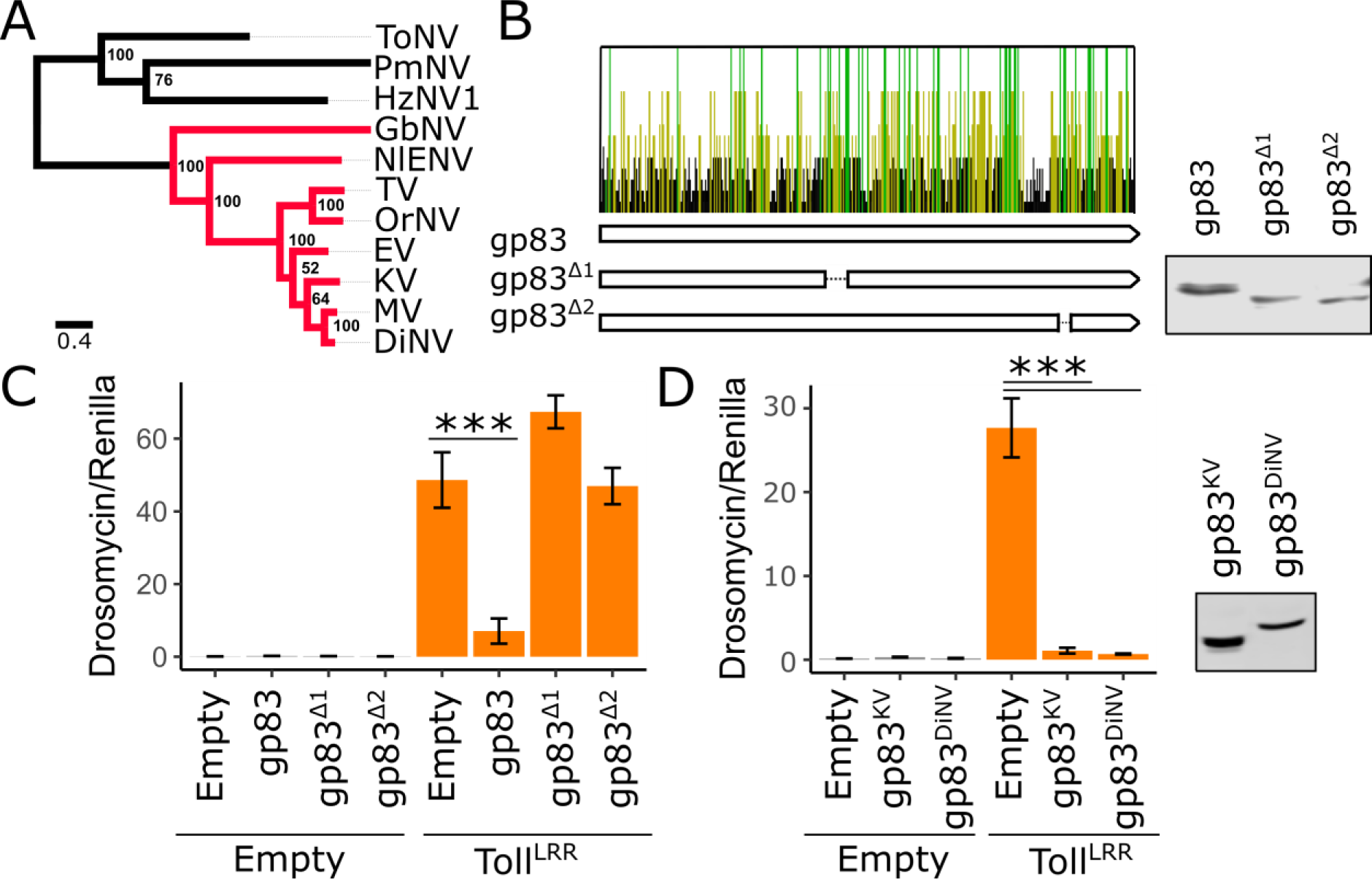
The immunosuppressive function of gp83 is evolutionarily conserved. (A) Maximum likelihood phylogeny inferred from a protein alignment of nudivirus-encoded DNA polymerase B using PhyML (Guindon *et al.*, 2010), with an LG substitution model and gamma-distributed rate parameter. Support for each node was assessed by bootstrapping, and the scale bar represents substitutions per site. Nudivirus species that encode gp83 homologs are coloured in red. (B)Conservation of gp83 amino acid residues across 7 species of nudivirus (all red viruses in panel A except the endogenized virus NlENV). Each bar represents an amino-acid residue, and bars are coloured yellow if the residue is conserved in ≥ 50% of the species shown in the phylogeny, and green if conserved in 100% of the species. Two deletion constructs were created which span regions with an excess of conserved residues: gp83^Δ1^ and gp83 ^Δ2^. Although protein accumulates to normal levels in both deletion constructs following transfection (B), overexpression of gp83 ^Δ1^ or gp83 ^Δ2^ failed to inhibit Toll^lrr^-induced Drs-FLuc expression (relative to pAct-FLuc expression). (C) Overexpression of gp83 from DiNV was able to inhibit Toll^lrr^-induced Drs-FLuc expression (relative to pAct-FLuc expression) in *D. melanogaster* cells, indicating the Toll suppressive function against *D. melanogaster* is conserved in other nudiviruses. Error bars show standard error of the mean (n = 5 biological replicates). *p < 0.05; **p < 0.01; ***p < 0.001 (Statistical tests performed in MCMCglmm)

## Conclusions

In this study we investigated the role of known anti-RNA viral immune pathways in the context of DNA virus infection, including RNAi, JAK-STAT, Imd, and Toll pathways. Our data support an antiviral role for RNAi and Imd against KV, consistent with previously-described antiviral RNAi against IIV6 and DNA virus-encoded suppressors of Imd (Bronkhorst *et al.*, 2012; Kemp *et al.*, 2013; Lamiable *et al.*, 2016). Furthermore, we identified gp83 as a KV-encoded Toll suppressor that acts downstream of NF-_k_B transcription factor release of ΙκΒ, strongly suggesting that Toll signalling can be antiviral during DNA virus infection in insects. The immunosuppressive effect of gp83 is conserved in other nudiviruses, and has not evolved host-specificity in DiNV, indicating that the Toll-gp83 interaction is unlikely to be a hotspot of reciprocal host-virus adaptation and that other KV genes may be more important in determining host range.

## Acknowledgements

We thank Maria-Carla Saleh, Marc Dionne, François Leulier, David Finnegan, and Bruno Lemaitre for kindly sharing RNAi, Toll, and Imd pathway mutant fly lines. We thank Pascale Dijkers, Jean-Luc Imler, Neal Silverman, Edan Foley, and Norbert Perrimon (Addgene plasmid # 37393) for kindly sharing Toll, Imd, and JAK-STAT constructs. We thank Rob Unckless for sharing a DiNV DNA sample with us. We thank the Ruth Steward and the Developmental Studies Hybridoma Bank for making the dorsal antibody available. WHP is supported by the Darwin Trust of Edinburgh and by an EMBO Short-Term Fellowship in RvR’s laboratory (Grant Number 7095). Work in RvR’s laboratory is supported by a European Research Council Consolidator Grant under the European Union’s Seventh Framework Programme (grant number ERC CoG 615680) and a VICI grant from the Netherlands Organization for Scientific Research (grant number 016.VICI.170.090).

---

**Figure S1:**
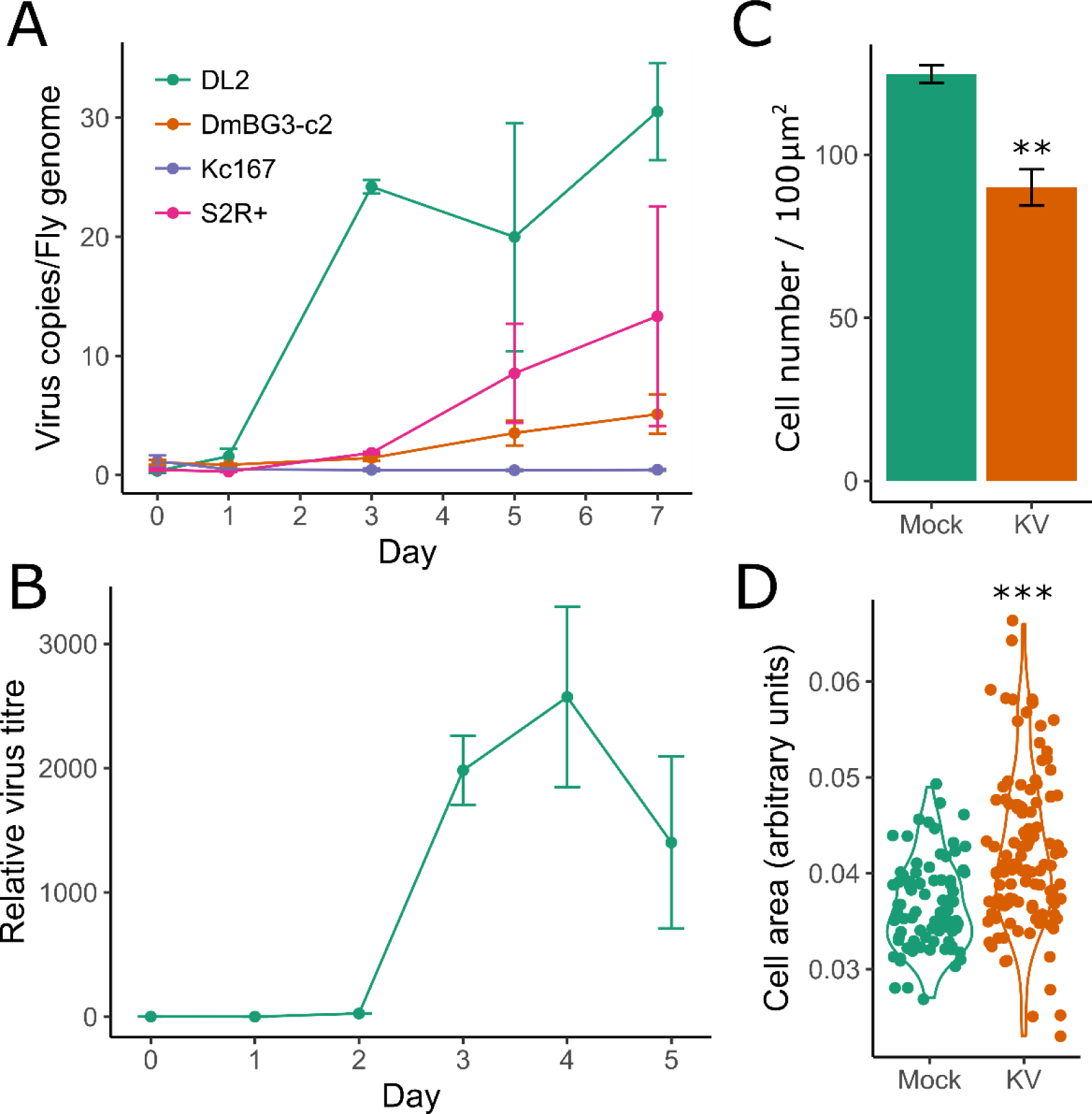
KV growth dynamics in cell culture. We assessed KV growth in various cell lines by qPCR for the KV genome, relative to the fly gene rpl32. KV had different growth rates across *D. melanogaster* cell lines (A), and grew best in DL2 and S2R+ cells (n=3 for each time point). (B) KV release from S2 cells into the culture medium was assessed by DNA extraction of 50 uL of culture medium and qPCR against the KV genome, plotted as relative to the amount of KV in the medium directly following infection (i.e. zero time point is equal to 1). KV was released from cells starting from 3 dpi (n = 3). Although we did not observe any overt cytopathic effects of KV-infected cells, passage of cells 7 dpi revealed a slower growth rate (C), evidenced by the number of cells per approximately 100 μm^2^ in KV versus mock-treated cells (n=3). (D) The passaged KV-infected cells also had a subset of significantly larger cells. Error bars show standard error of the mean. *p < 0.05; **p < 0.01; ***p < 0.001 (Statistical tests performed in MCMCglmm)

**Figure S2:**
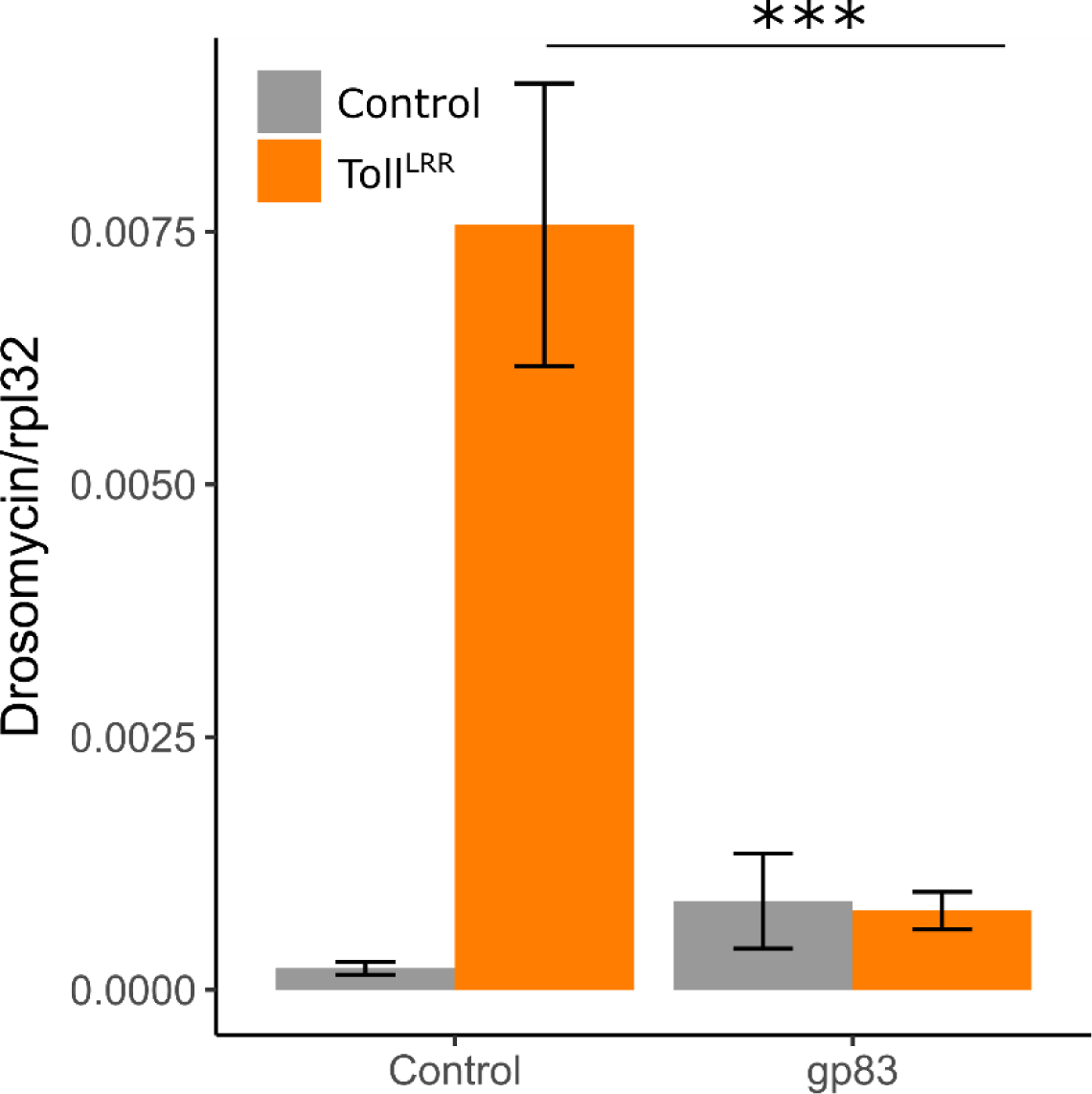
gp83 Inhibits endogenous Drosomycin expression. We performed qRT-PCR of Drs, measured relative to rpl32 expression, confirming that gp83 is also able to inhibit the expression of endogenously encoded AMPs. Error bars show standard error of the mean (n = 5 biological replicates). ***p < 0.001 (Statistical tests performed in MCMCglmm)

**Figure S3:**
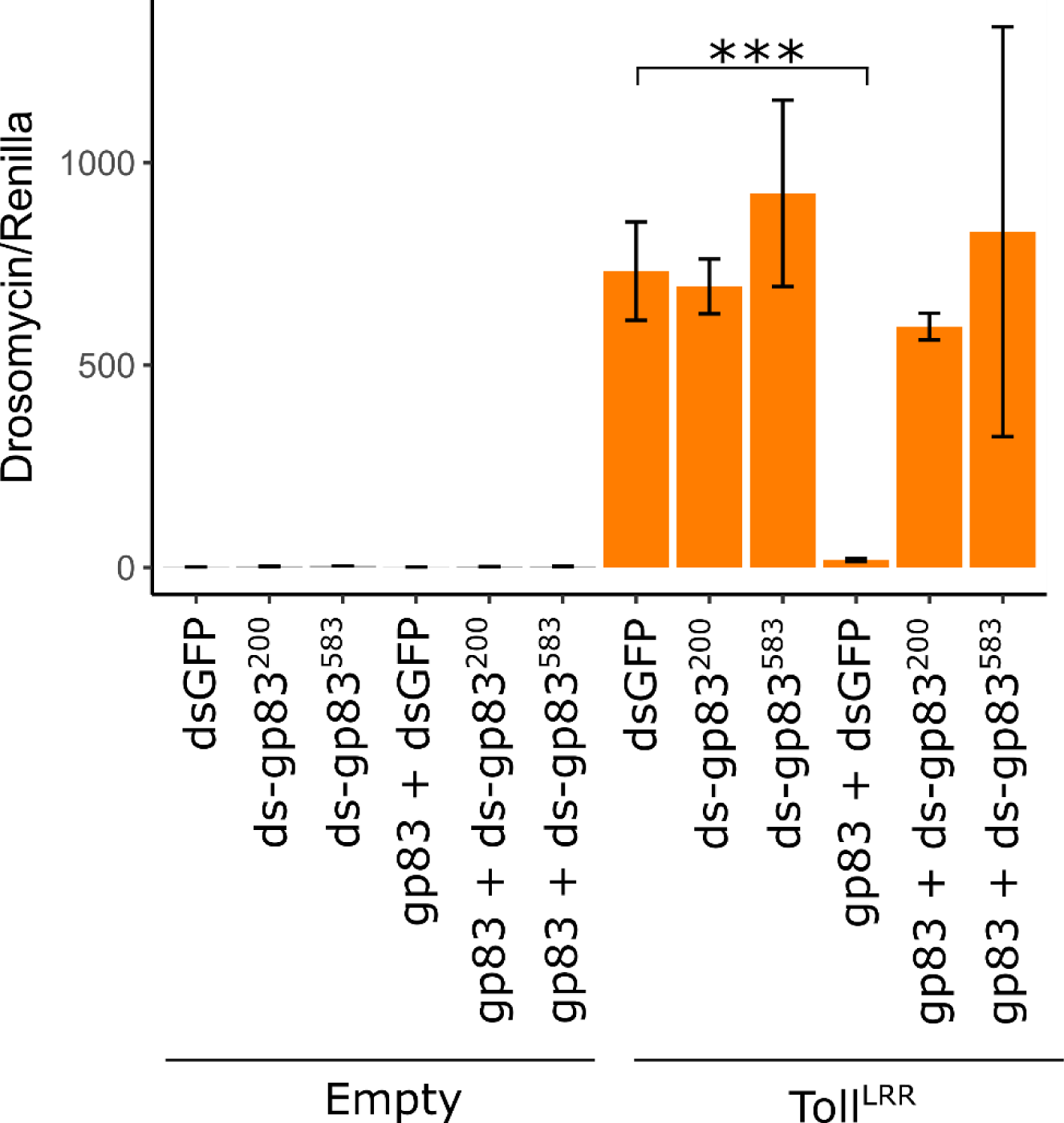
Confirmation of gp83 knock down efficiency. Co-transfection of gp83 with two independent dsRNAs against gp83 was able to reverse the immunosuppressive effect of gp83 on Toll signalling, measured by Drs-FLuc expression (relative to pAct-FLuc expression), confirming these dsRNAs were efficient in knocking down gp83 expression. Error bars show standard error of the mean (n = 3 biological replicates). ***p < 0.001 (Statistical tests performed in MCMCglmm)

**Figure S4:**
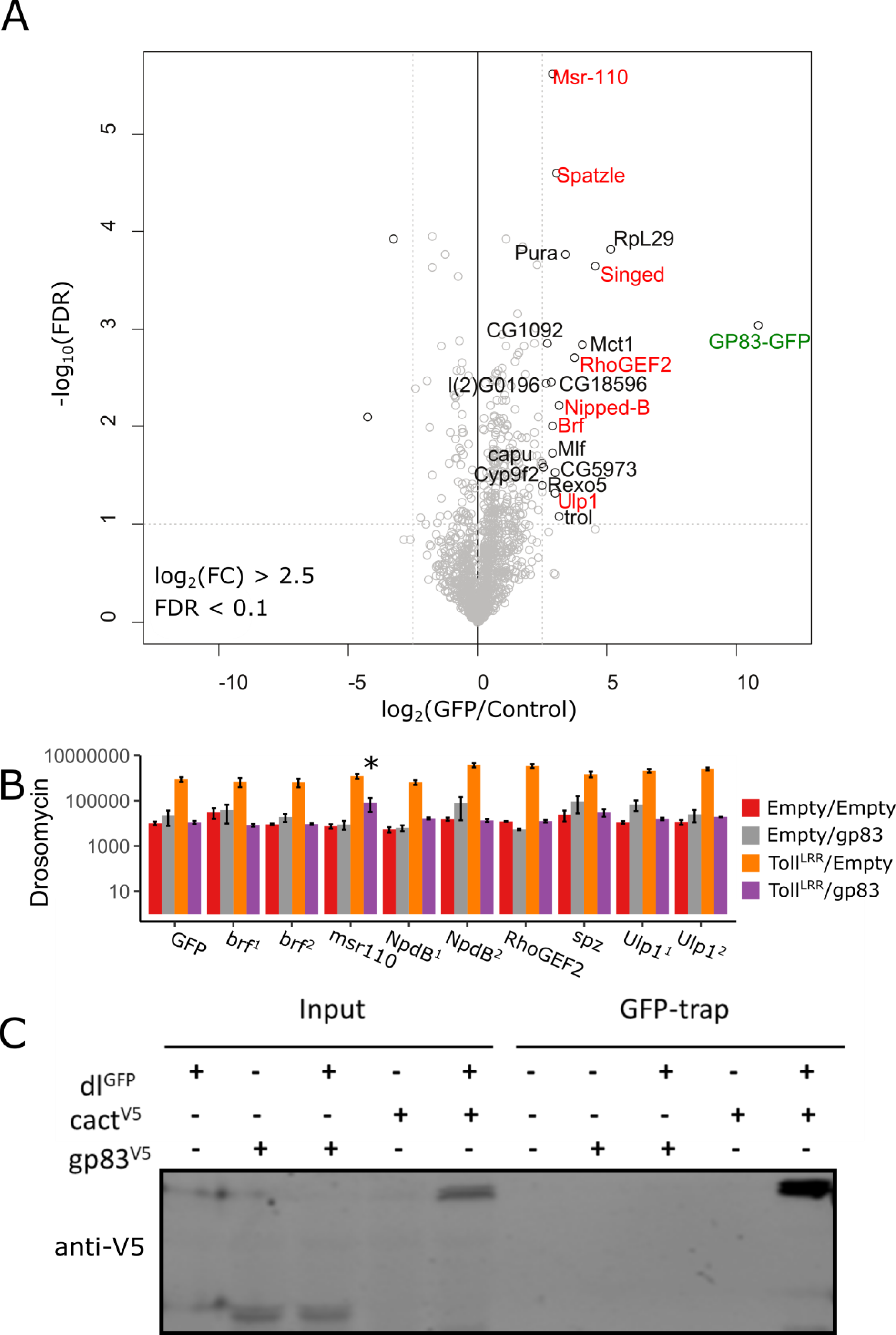
Identification of host interactors of gp83. (A) Identification of gp83 interacting proteins in S2 cell lysates by label-free quantitative (LFQ) mass spectrometry. Permutation-based FDR-corrected t-tests were used to determine proteins that are statistically enriched in gp83-GFP IP. The log2 LFQ intensity of GFP-gp83 IP over control IP (cells that do not express gp83-GFP) is plotted against the –log_10_ FDR. Interactors with an enrichment of fold change > 2.5; -log_10_ FDR > 1 are indicated. Although the bait, gp83-GFP (labelled in green), was efficiently retained, few proteins were strongly enriched. (B) Through dsRNA knock-down, we confirmed brf, msr-110, Nipped-B, RhoGEF2, spatzle, and Ulp1 (shown labelled in red in panel A) are not involved in Toll signalling, or gp83 suppression of Toll signalling, as measured by Drs-FLuc expression (relative to pAct-FLuc expression). Genes are superscripted with ‘1’ or ‘2’ when two independent dsRNAs were used to knockdown the gene. Although msr-110 appears to partially rescue gp83 immunosuppression, subsequent experiments did not reproduce this effect. Error bars represent standard error of the mean (n=3). Statistical tests were performed in MCMCglmm. (C) We were unable to identify an association between gp83^v5^ and dl^gfp^ through dl^gfp^ immunoprecipitation. We used the previously described interaction between cact (V5 tagged) as a positive control.

**Figure S5:**
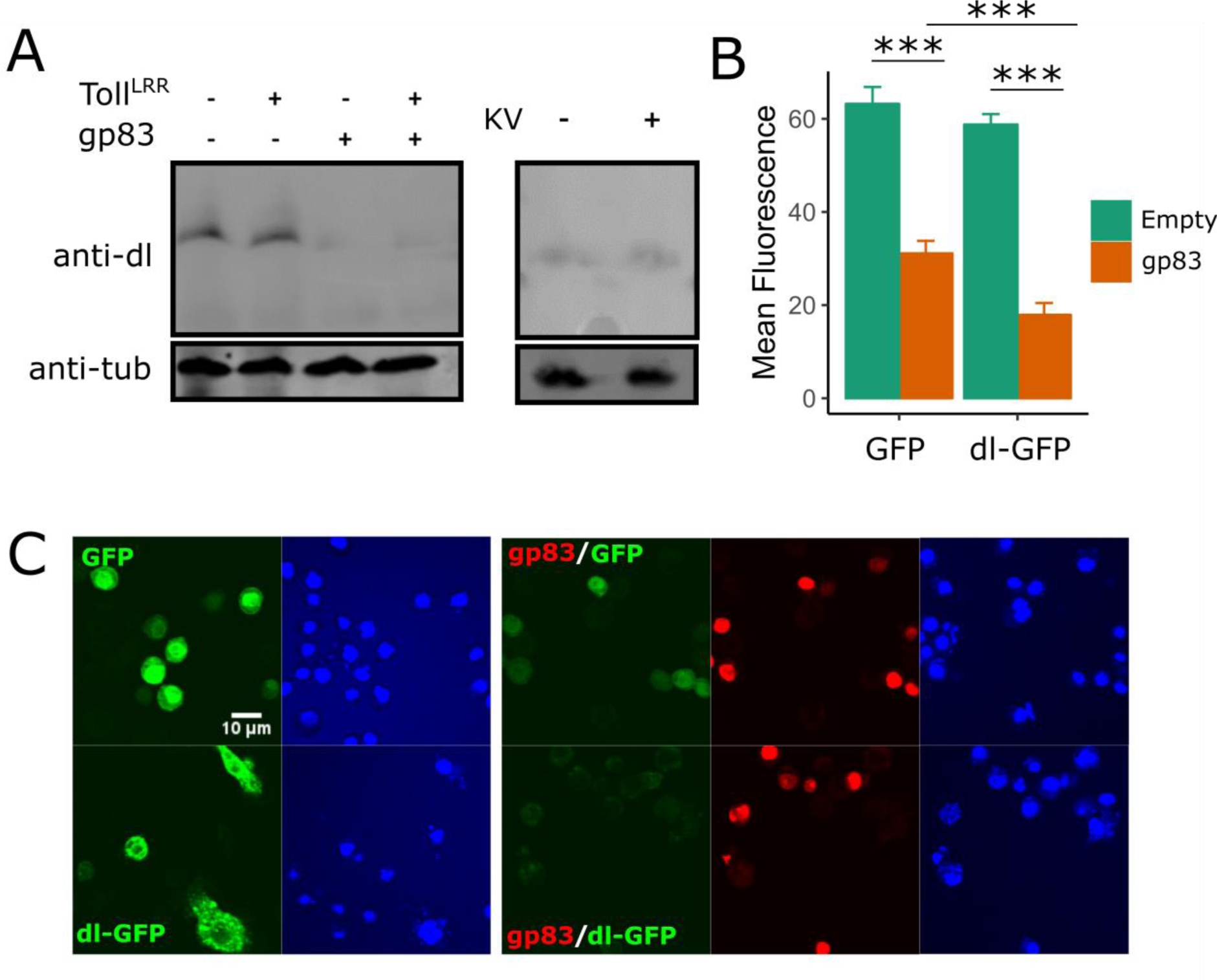
Overexpression of gp83 may reduce dorsal levels. (A) Overexpression of gp83 led to reduced levels of endogenous dl observed on western blot, regardless of Toll activation (i.e. Toll^lrr^ overexpression; left panel). However, KV infection did not reduce dl staining, indicating dl degradation may not be the primary mechanism by which KV inhibits Toll signalling (A, right). (B) ImageJ was used to quantify mean GFP fluorescence for individually outlined cells transfected with either GFP or dl-GFP, with or without gp83 (n ≥ 20 cells for each condition). Overexpression of gp83 inhibited both GFP and dl-GFP levels, although it reduced dl-GFP levels by significantly more (MCMCp < 0.001). Error bars show standard error of the mean. (C) A representative image from (B), showing GFP and dl-GFP expression with or without gp83 overexpression ***p < 0.001

